# Creating a universal SNP and small indel variant caller with deep neural networks

**DOI:** 10.1101/092890

**Authors:** Ryan Poplin, Pi-Chuan Chang, David Alexander, Scott Schwartz, Thomas Colthurst, Alexander Ku, Dan Newburger, Jojo Dijamco, Nam Nguyen, Pegah T. Afshar, Sam S. Gross, Lizzie Dorfman, Cory Y. McLean, Mark A. DePristo

## Abstract

Next-generation sequencing (NGS) is a rapidly evolving set of technologies that can be used to determine the sequence of an individual’s genome^1^ by calling genetic variants present in an individual using billions of short, errorful sequence reads^2^. Despite more than a decade of effort and thousands of dedicated researchers, the hand-crafted and parameterized statistical models used for variant calling still produce thousands of errors and missed variants in each genome^3,4^. Here we show that a deep convolutional neural network^5^ can call genetic variation in aligned next-generation sequencing read data by learning statistical relationships (likelihoods) between images of read pileups around putative variant sites and ground-truth genotype calls. This approach, called DeepVariant, outperforms existing tools, even winning the “highest performance” award for SNPs in a FDA-administered variant calling challenge. The learned model generalizes across genome builds and even to other mammalian species, allowing non-human sequencing projects to benefit from the wealth of human ground truth data. We further show that, unlike existing tools which perform well on only a specific technology, DeepVariant can learn to call variants in a variety of sequencing technologies and experimental designs, from deep whole genomes from 10X Genomics to Ion Ampliseq exomes. DeepVariant represents a significant step from expert-driven statistical modeling towards more automatic deep learning approaches for developing software to interpret biological instrumentation data.

## Main Text

Calling genetic variants from NGS data has proven challenging because NGS reads are not only errorful (with rates from ∼0.1-10%) but result from a complex error process that depends on properties of the instrument, preceding data processing tools, and the genome sequence itself^1,3,4,6^. State-of-the-art variant callers use a variety of statistical techniques to model these error processes and thereby accurately identify differences between the reads and the reference genome caused by real genetic variants and those arising from errors in the reads^3,4,6,7^. For example, the widely-used GATK uses logistic regression to model base errors, hidden Markov models to compute read likelihoods, and naive Bayes classification to identify variants, which are then filtered to remove likely false positives using a Gaussian mixture model with hand-crafted features capturing common error modes^6^. These techniques allow the GATK to achieve high but still imperfect accuracy on the Illumina sequencing platform^3,4^. Generalizing these models to other sequencing technologies has proven difficult due to the need for manual retuning or extending these statistical models (see e.g. Ion Torrent^8,9^), a major problem in an area with such rapid technological progress^1^.

Here we describe a variant caller for NGS data that replaces the assortment of statistical modeling components with a single, deep learning model. Deep learning is a machine learning technique applicable to a variety of domains, including image classification^10^, translation^11^, gaming^12,13^, and the life sciences^14–17^. This toolchain, which we call DeepVariant, (Figure 1) begins by finding candidate SNPs and indels in reads aligned to the reference genome with high-sensitivity but low specificity. The deep learning model, using the Inception-v2 architecture^5^, emits probabilities for each of the three diploid genotypes at a locus using a pileup image of the reference and read data around each candidate variant (Figure 1). The model is trained using labeled true genotypes, after which it is frozen and can then be applied to novel sites or samples. Throughout the following experiments, DeepVariant was trained on an independent set of samples or variants to those being evaluated.

**Figure 1:**
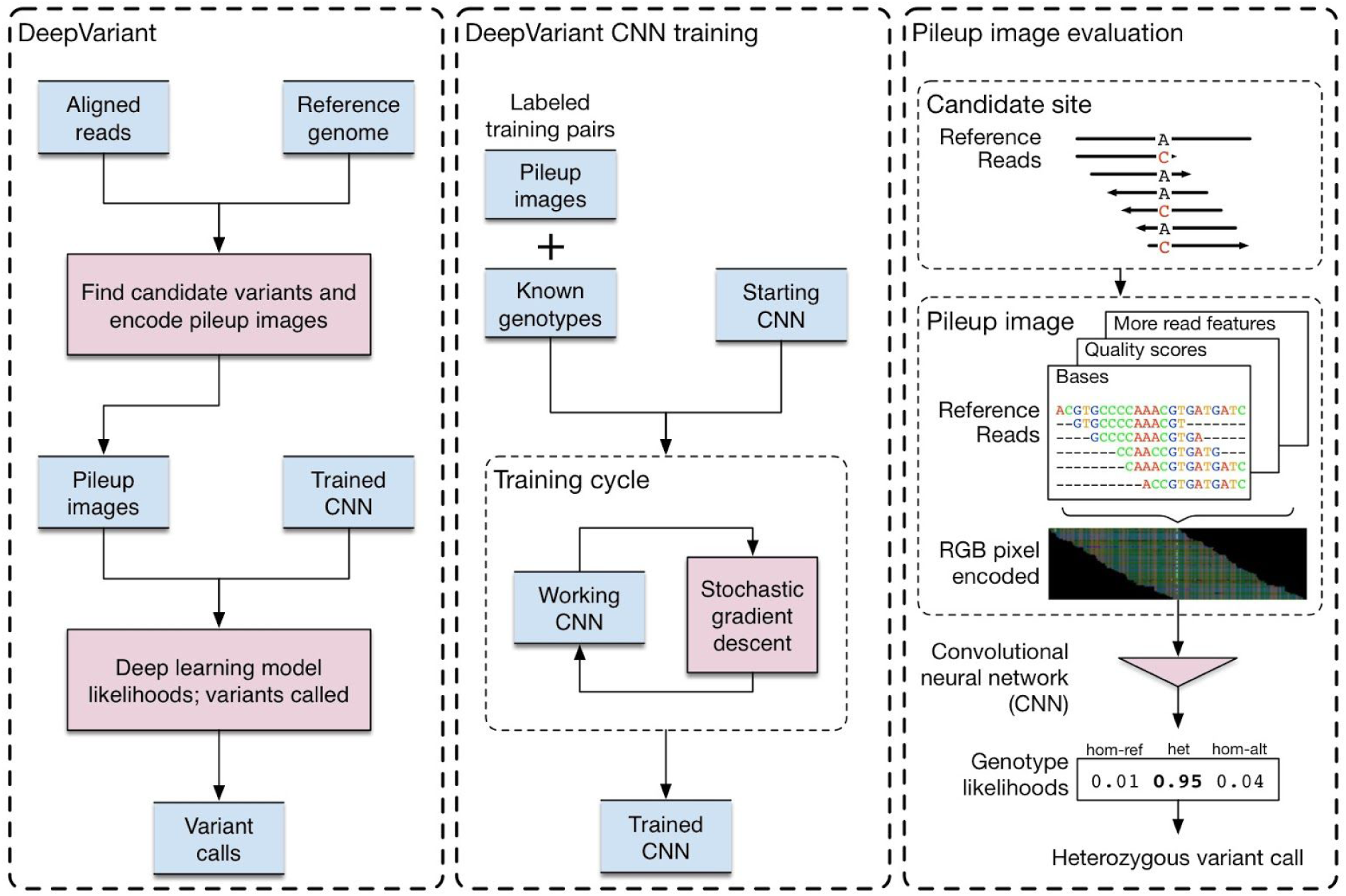
DeepVariant workflow overview. Before DeepVariant, NGS reads are first aligned to a reference genome and cleaned up with duplicate marking and, optionally, local assembly. (Left box) First, the aligned reads are scanned for sites that may be different from the reference genome. The read and reference data is encoded as an image for each candidate variant site. A trained convolutional neural network (CNN) calculates the genotype likelihoods for each site. A variant call is emitted if the most likely genotype is heterozygous or homozygous non-reference. (Middle box) Training the CNN reuses the DeepVariant machinery to generate pileup images for a sample with known genotypes. These labeled image + genotype pairs, along with an initial CNN which can be a random model, a CNN trained for other image classification tests, or a prior DeepVariant model, are used to optimize the CNN parameters to maximize genotype prediction accuracy using a stochastic gradient descent algorithm. After a maximum number of cycles or time has elapsed or the model’s performance has convergence, the final trained model is frozen and can then be used for variant calling. (Right box) The reference and read bases, qualities scores, and other read features are encoded into an red-green-blue (RGB) pileup image at a candidate variant. This encoded image is provided to the CNN to calculate of the genotype likelihoods for the three diploid genotype states of homozygous reference (hom-ref), heterozygous (het), or homozygous alternate (hom-alt). In this example a heterozygous variant call is emitted as the most probable genotype likelihood here is “het”. In all panels blue boxes represent data and red boxes are processes. Details of all processes are given in the “Online methods”.

This deep learning model was trained without specialized knowledge about genomics or next-generation sequencing, and yet can learn to call genetic variants more accurately than state-of-the-art methods. When applied to the Platinum Genomes Project NA12878 data^18^, DeepVariant produces a callset with better performance than the GATK when evaluated on the held-out chromosomes of the Genome in a Bottle ground truth set (Figure 2A, Figure S1). For further validation, we sequenced 35 replicates of NA12878 using a standard whole-genome sequencing protocol and called variants on 27 replicates using a GATK best-practices pipeline and DeepVariant using a model trained on the other eight replicates (see methods). Not only does DeepVariant produce more accurate results but it does so with greater consistency across a variety of quality metrics (Figure 2B).

**Figure 2:**
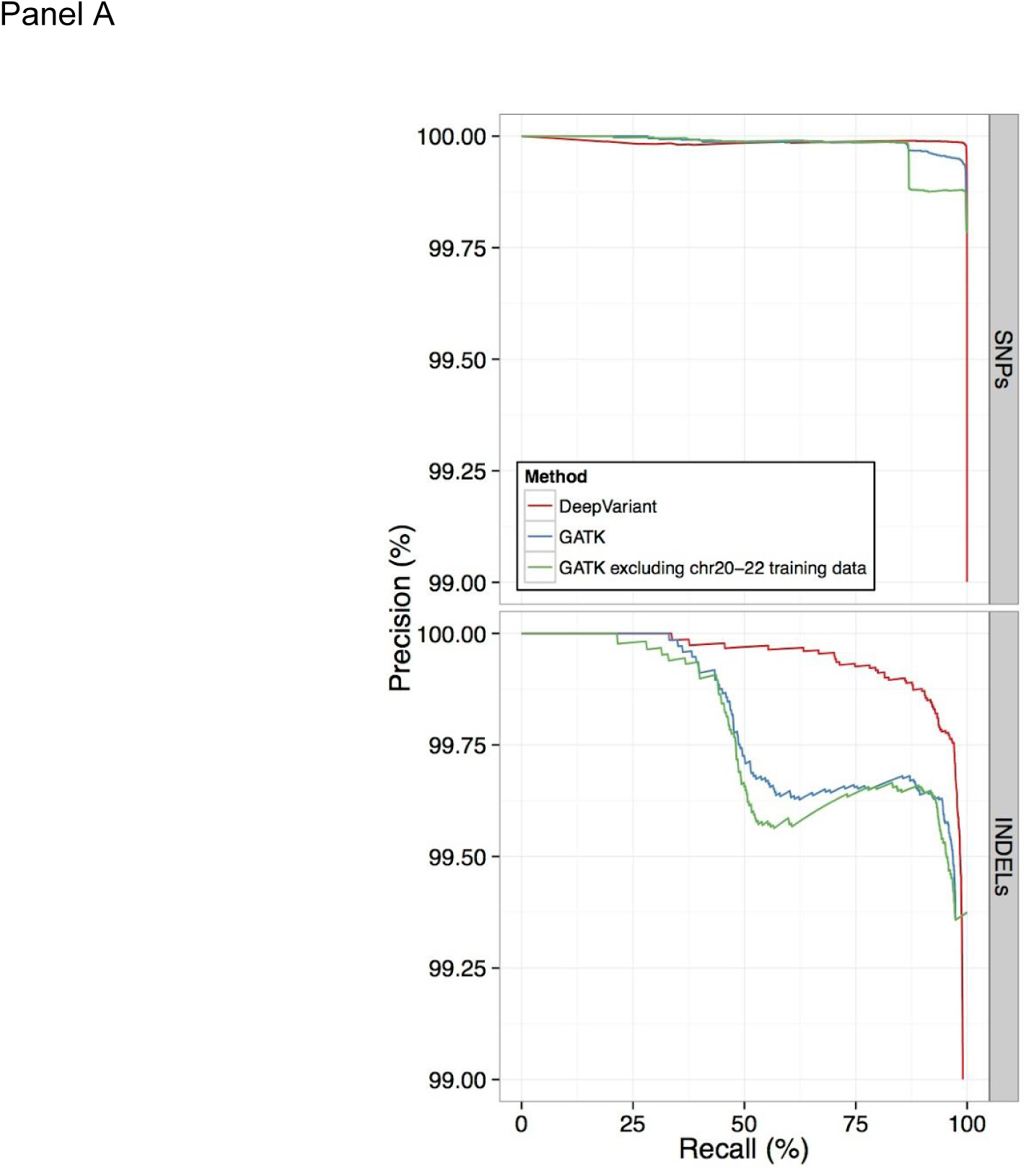
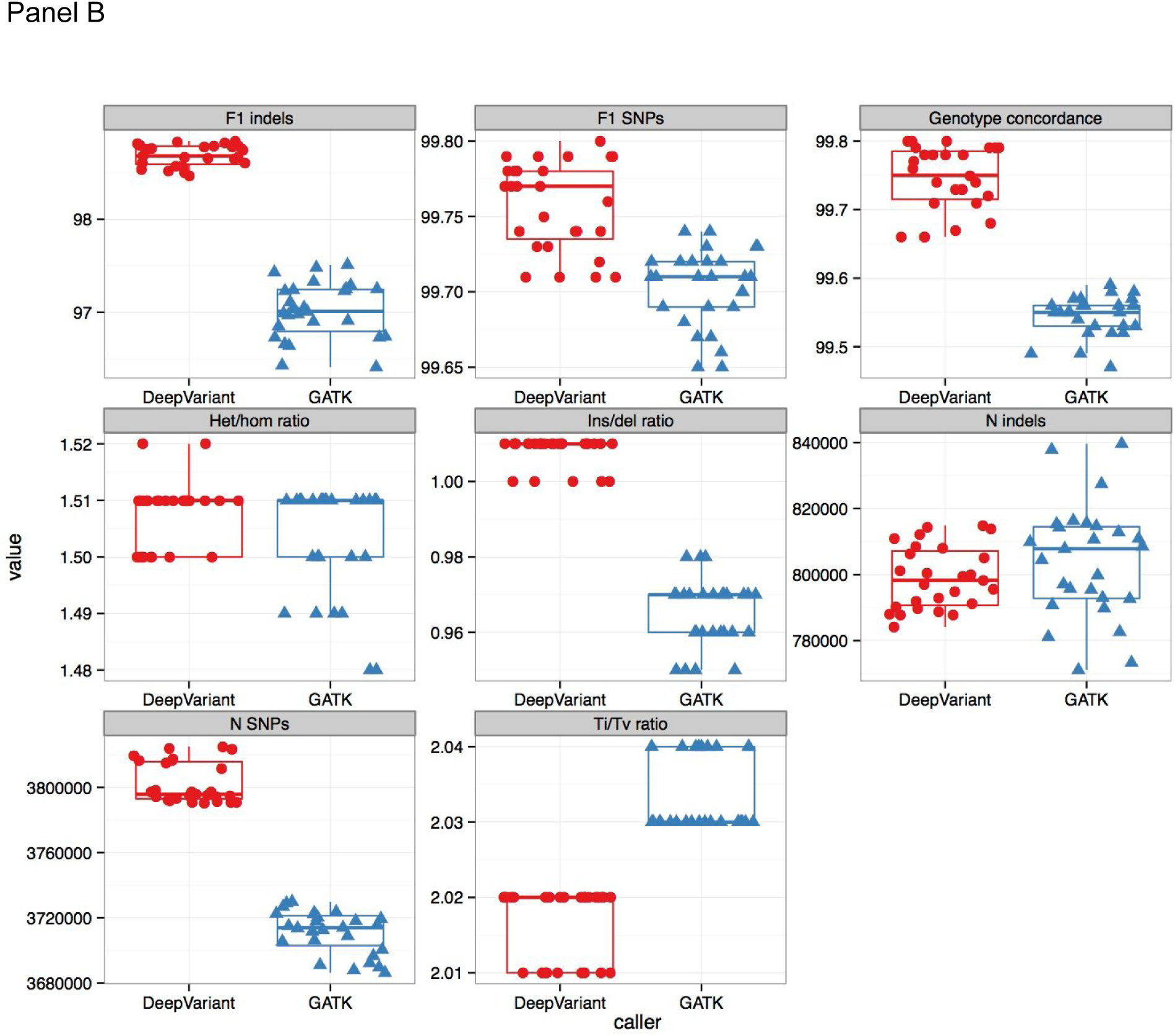
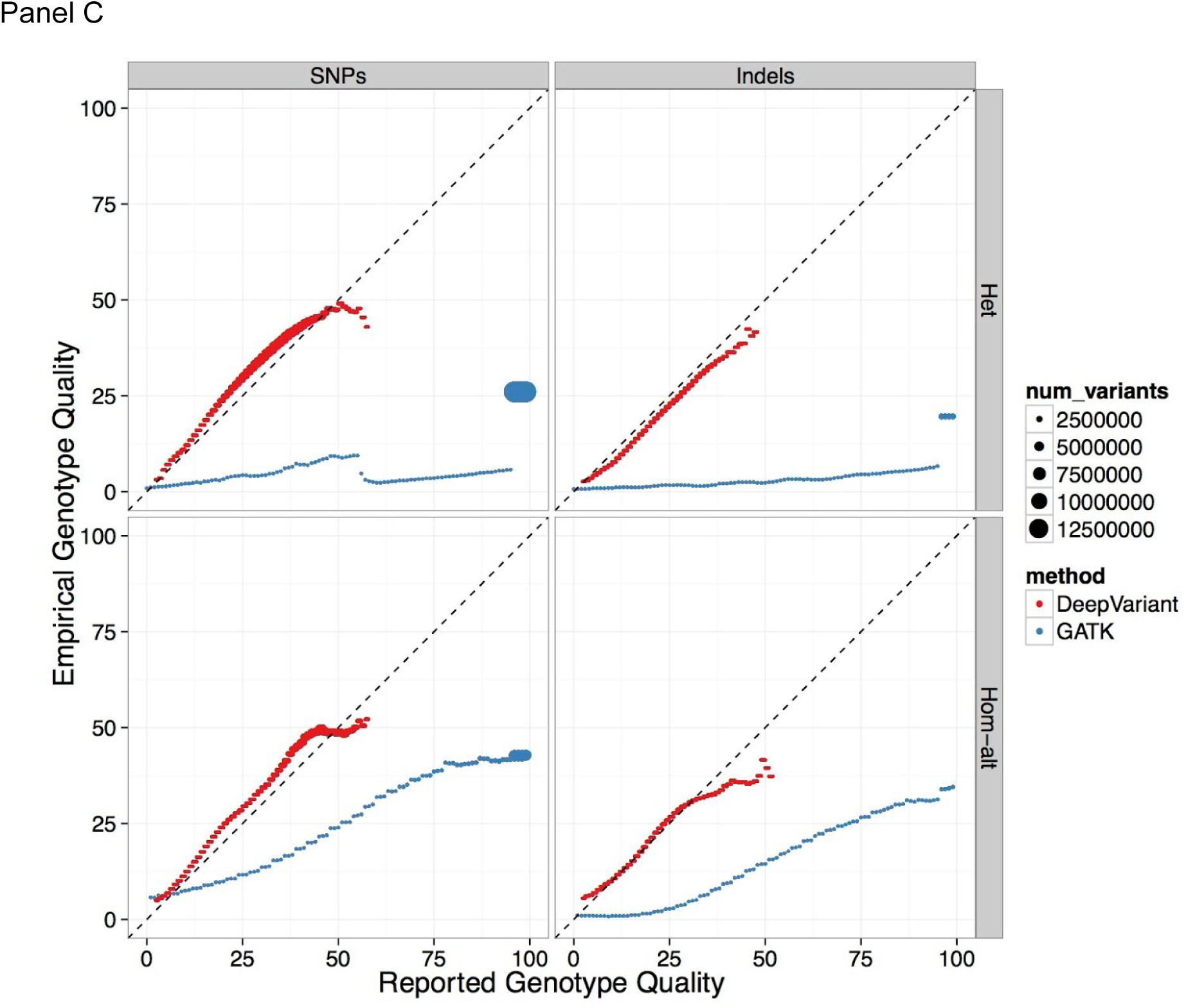
DeepVariant accuracy, consistency, and calibration relative to the GATK. (A) Precision-recall plot for DeepVariant (red) and GATK (green, blue) calls for the Genome in the Bottle benchmark sample NA12878 using 2×101 Illumina HiSeq data from the Platinum Genomes project. The GATK was run in two ways. In the first, GATK best-practices were followed and the variant filtering step (VQSR) was provided data for known variants on both the training and test chromosomes, allowing VQSR to use population variation information to better call variants on the test chromosomes. In the second, we removed all population variation information for the test chromosomes chr20-22, relying on the VQSR model learned only on the training chromosomes, which is more representative of the GATK’s calling performance on novel variation. See Supp. Mats. for additional details and figures. (B) DeepVariant (red circles) and the GATK (blue triangles) were run on 27 independently sequenced replicates of NA12878 (PCR-free WGS 2×151 on an Illumina X10 with coverage from 24×-35×). Each panel shows the distribution of values for the given metric (panel label) for DeepVariant and the GATK. DeepVariant produces more accurate SNP and indel calls (F1) when compared to the Genome in a Bottle standard for NA12878 with a higher fraction of sites having the correct genotype assigned (Genotype concordance). DeepVariant finds a similar numbers of indels to the GATK, but has a more consistent ratio of insertions to deletions. DeepVariant finds more SNPs than GATK with a similar ratio of heterozygous variants to homozygous alternative variants (Het/hom ratio). (C) Comparison of likelihoods assigned to heterozygous and homozygous alternate genotypes emitted by DeepVariant and the GATK shows the likelihood model learned by DeepVariant is substantially better calibrated than that employed by the GATK. On the x-axis is the reported genotype quality (GQ) for calls for DeepVariant (red) and GATK (blue) compared to the observed error rate in each of these GQ bands (y-axis), for true heterozygous and homozygous variants (vertical facet) and SNPs and indels (horizontal facet) separately. The size of each calibration point reflects the number of variant calls used to estimate the empirical accuracy. The calibration curves were calculated using genotype likelihoods from the held-out evaluation data in eight sequenced replicates of NA12878. For example, the set of all Q30 heterozygous calls should be in aggregate accurate at a rate of 999 in 1000. Genotypes should be correct at a rate declared by their confidence; perfect calibration would follow the marked x=y line.

Like many variant calling algorithms, the GATK relies on a model that assumes read errors are independent^6^. Though long-recognized as an invalid assumption^2^, the true likelihood function that models multiple reads simultaneously is unknown^6,19,20^. Because DeepVariant presents an image of all of the reads relevant for a putative variant together, the convolutional neural network (CNN) is able to account for the complex dependence among the reads by virtue of being a universal approximator^21^. This manifests itself as a tight concordance between the estimated probability of error from the likelihood function and the observed error rate, as seen in Figure 2C where DeepVariant’s CNN is well calibrated, strikingly more so than the GATK. That the CNN has approximated this true, but unknown, inter-dependent likelihood function is the essential technical advance enabling us to replace the hand-crafted statistical models in other approaches with a single deep learning model and still achieve such high performance in variant calling.

To further confirm the performance of DeepVariant, we submitted variant calls for a blinded sample, NA24385, to the Food and Drug Administration-sponsored variant calling <u>Truth Challenge</u> in May 2016 and won the “highest performance” award for SNPs by an independent team using a different evaluation methodology. For this contest DeepVariant was trained only on data available from the CEPH female sample NA12878 and was evaluated on the unseen Ashkenazi male sample NA24385, achieving high accuracy (SNP F1 = 99.95%, indel F1 = 98.98%) and showing that DeepVariant can generalize beyond its training data. We then applied the same dataset and evaluation methodology to a variety of both recent and commonly used bioinformatics methods including the GATK, FreeBayes^22^, samtools^23^, 16GT^24^, and Strelka^25^ (Table 1). DeepVariant demonstrated a more than 50% reduction in total number of errors per genome (4,652 errors) compared to the next best algorithm (9,531 errors). We also evaluated the same set of methods using the synthetic diploid sample CHM1-CHM13^26^ (Table 2). In our tests DeepVariant outperformed all other methods for calling both SNP and indel mutations without needing to adjust filtering thresholds or other parameters.

**Table 1:**
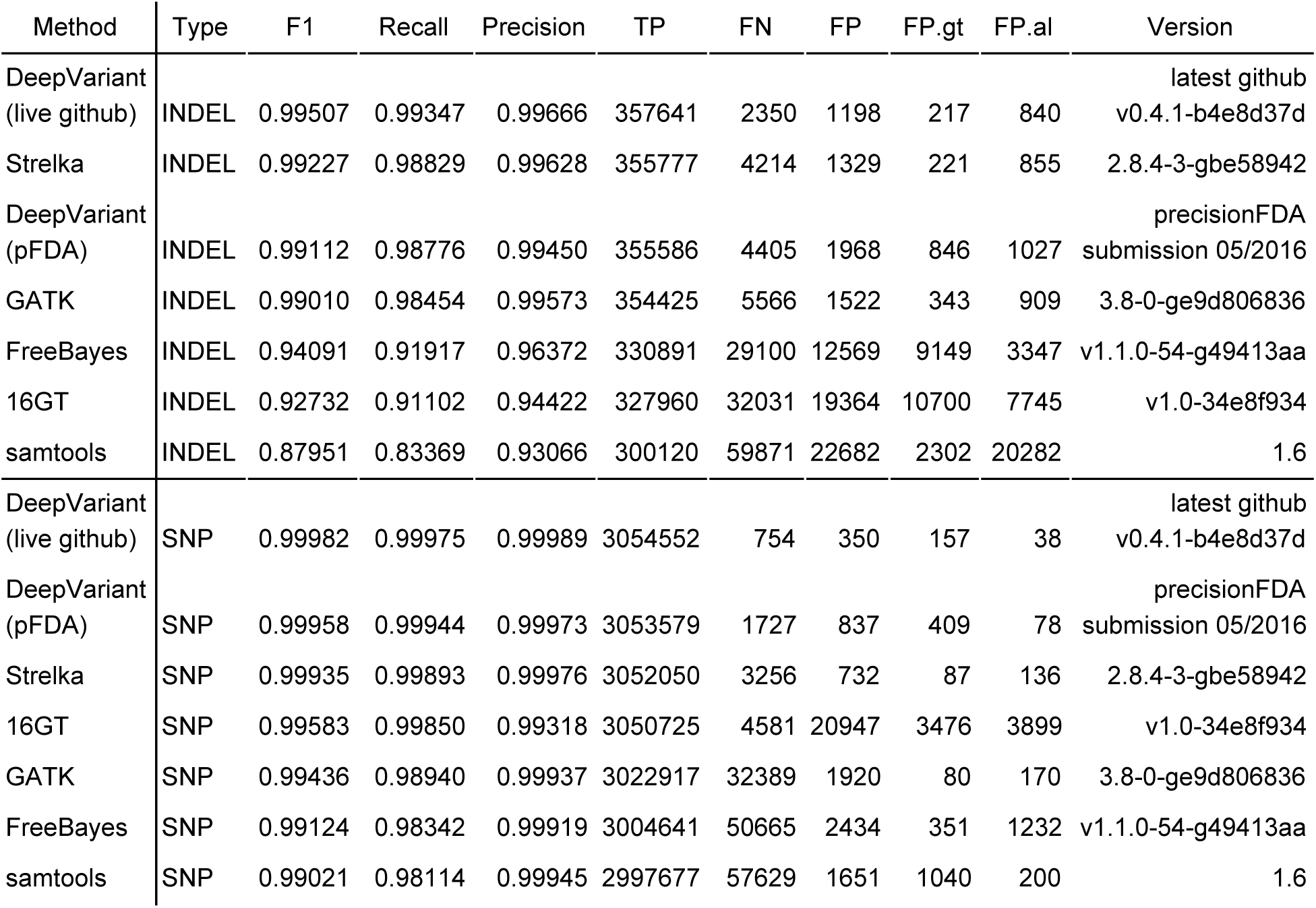
Evaluation of several bioinformatics methods on the high coverage, whole genome sample NA24385. The dataset used in this evaluation is the same as in the precisionFDA Truth Challenge. Several methods are compared including the DeepVariant callset as submitted to the contest as well as the most recent DeepVariant version from GitHub. Each method was run according to the individual authors’ best practice recommendations and represents a good faith effort to achieve best results. Comparisons to the Genome in a Bottle truth set for this sample was performed using the hap.py software available on GitHub at http://github.com/Illumina/hap.py using the same version of the GIAB truth set (v3.2.2) used by precisionFDA. The overall accuracy (F1, sort order within each variant type), recall, precision, and numbers of true positives (TP), false negatives (FN), and false positives (FP) are shown over the whole genome. False positives are further divided by those caused by genotype mismatches (FP.gt) and those cause by allele mismatches (FP.al). Finally, the version of the software used for each method is provided.

**Table 2:** Evaluation of several bioinformatics methods on the high coverage, whole genome synthetic diploid sample CHM1-CHM13. Several methods are compared including the most recent DeepVariant version from GitHub. Each method was run according to the individual authors’ best practice recommendations and represents a good faith effort to achieve best results. Comparisons to the CHM1-CHM13 truth set was performed using the CHM-eval.kit software available on GitHub at https://github.com/lh3/CHM-eval release version 0.5. The overall accuracy (F1, sort order within each variant type), recall, precision, and numbers of true positives (TP), false negatives (FN), and false positives (FP) are shown over the whole genome. Finally, the version of the software used for each method is provided.

| Method | Type | F1 | Recall | Precision | TP | FN | FP | Version |
| --- | --- | --- | --- | --- | --- | --- | --- | --- |
| DeepVariant | INDEL | 0.95806 | 0.92868 | 0.98936 | 529137 | 40634 | 5690 | v0.4.1-b4e8d37d |
| Strelka | INDEL | 0.95074 | 0.91623 | 0.98796 | 522039 | 47732 | 6363 | 2.8.4-3-gbe58942 |
| 16GT | INDEL | 0.94010 | 0.90803 | 0.97452 | 517369 | 52402 | 13527 | v1.0-34e8f934 |
| GATK | INDEL | 0.91212 | 0.84497 | 0.99087 | 481441 | 88330 | 4437 | 3.8-0-ge9d806836 |
| FreeBayes | INDEL | 0.90438 | 0.83025 | 0.99305 | 473053 | 96718 | 3313 | v1.1.0-54-g49413aa |
| Samtools | INDEL | 0.86976 | 0.79089 | 0.96611 | 450626 | 119145 | 15807 | 1.6 |
| DeepVariant | SNP | 0.99103 | 0.98888 | 0.99319 | 3518118 | 39553 | 24132 | v0.4.1-b4e8d37d |
| Strelka | SNP | 0.98865 | 0.98107 | 0.99636 | 3490314 | 67357 | 12749 | 2.8.4-3-gbe58942 |
| 16GT | SNP | 0.97862 | 0.98966 | 0.96782 | 3520894 | 36777 | 117078 | v1.0-34e8f934 |
| FreeBayes | SNP | 0.96910 | 0.94837 | 0.99075 | 3373984 | 183687 | 31492 | v1.1.0-54-g49413aa |
| GATK | SNP | 0.96895 | 0.94542 | 0.99368 | 3363476 | 194195 | 21379 | 3.8-0-ge9d806836 |
| Samtools | SNP | 0.96818 | 0.94386 | 0.99378 | 3357947 | 199724 | 21012 | 1.6 |

We further explored how well DeepVariant’s CNN generalizes beyond its training data. First, a model trained with read data aligned to human genome build GRCh37 and applied to reads aligned to GRCh38 has similar performance (overall F1 = 99.45%) to one trained on GRCh38 and then applied to GRCh38 (overall F1 = 99.53%), thereby demonstrating that a model learned from one version of the human genome reference can be applied to other versions with effectively no loss in accuracy (Table S1). Second, models learned using human reads and ground truth data achieve high accuracy when applied to a mouse dataset^27^ (F1 = 98.29%), out-performing training on the mouse data itself (F1 = 97.84%, Table S4). This last experiment is especially demanding as not only do the species differ but nearly all of the sequencing parameters do as well: 50× 2×148bp from an Illumina TruSeq prep sequenced on a HiSeq 2500 for the human sample and 27× 2×100bp reads from a custom sequencing preparation run on an Illumina Genome Analyzer II for mouse^27^. Thus, DeepVariant is robust to changes in sequencing depth, preparation protocol, instrument type, genome build, and even mammalian species. The practical benefits of this capability is substantial, as DeepVariant enables resequencing projects in non-human species, which often have no ground truth data to guide their efforts^27,28^, to leverage the large and growing ground truth data in humans.

To further assess its capabilities, we trained DeepVariant to call variants in eight datasets from Genome in a Bottle^29^ that span a variety of sequencing instruments and protocols, including whole genome and exome sequencing technologies, with read lengths from fifty to many thousands of basepairs (Table 3 and S6). We used the already processed BAM files to introduce additional variability as these BAMs differ in their alignment and cleaning steps. The results of this experiment all exhibit a characteristic pattern: the candidate variants have the highest sensitivity but a low PPV (mean 57.6%), which varies significantly by dataset. After retraining, all of the callsets achieve high PPVs (mean of 99.3%) while largely preserving the candidate callset sensitivity (mean loss of 2.3%). The high PPVs and low loss of sensitivity indicate that DeepVariant can learn a model that captures the technology-specific error processes in sufficient detail to separate real variation from false positives with high fidelity for many different sequencing technologies.

**Table 3:** Training DeepVariant to call variants on a variety of sequencing technologies and experimental protocols. Datasets are labeled to indicate instrument, protocol, target area (WGS for whole genome, gene regions as exome), with sequencing depth shown for whole genome targets. For each dataset, a set of candidate variants were identified across the genome in the NGS reads (methods). The baseline Illumina model was retrained using the candidate variants with labeled genotypes on chromosomes 1-19. This retrained model was then used to assign genotype likelihoods to the candidate variants, keeping those confidently non-reference on the held-out chromosomes 20-22. The sensitivity, positive predictive value (PPV), and overall accuracy (F1) are shown for the candidate and called variants on chr20-22 only.

| Dataset | Sensitivity |  | PPV |  | F1 |  |
| --- | --- | --- | --- | --- | --- | --- |
|  | Candidate variants | Called variants | Candidate variants | Called variants | Candidate variants | Called variants |
| 10X Chromium 75x WGS | 99.66% | 98.73% | 89.55% | 99.91% | 94.34% | 99.32% |
| 10X GemCode 34x WGS | 97.03% | 94.34% | 75.19% | 99.47% | 84.73% | 96.84% |
| Illumina HiSeq 31x WGS | 99.88% | 99.76% | 95.14% | 99.98% | 97.45% | 99.87% |
| Illumina HiSeq 60x WGS | 99.95% | 99.88% | 90.90% | 99.98% | 95.21% | 99.93% |
| Ion AmpliSeq exome | 91.94% | 89.28% | 8.05% | 99.70% | 14.81% | 94.21% |
| PacBio 40x WGS | 93.36% | 88.51% | 22.14% | 97.25% | 35.79% | 92.67% |
| SOLID SE 85x WGS | 82.50% | 76.62% | 14.27% | 99.01% | 24.33% | 86.39% |
| Illumina TruSeq exome | 94.04% | 92.58% | 65.31% | 99.31% | 77.08% | 95.83% |
| Mean | 94.79% | 92.46% | 57.57% | 99.33% | 65.47% | 95.63% |
| Median | 94.79% | 92.58% | 65.31% | 99.47% | 77.08% | 95.83% |

As we already shown above that DeepVariant performs well on Illumina WGS data, we analyze here the behavior of DeepVariant on two non-Illumina WGS datasets and two exome datasets from Illumina and Ion Torrent. The SOLID and Pacific Biosciences (PacBio) WGS datasets have high error rates in the candidate callsets. SOLID (13.9% PPV for SNPs, 96.2% for indels, and 14.3% overall) has many SNP artifacts from the mapping short, color-space reads. The PacBio dataset is the opposite, with many false indels (79.8% PPV for SNPs, 1.4% for indels, and 22.1% overall) due to this technology’s high indel error rate. Training DeepVariant to call variants in an exome is likely to be particularly challenging. Exomes have far fewer variants (∼20-30k)^30^ than found in a whole-genome (∼4-5M)^31^. The non-uniform coverage and sequencing errors from the exome capture or amplification technology also introduce many false positive variants^32^. For example, at 8.1% the PPV of our candidate variants for Ion Ampliseq is the lowest of all our datasets.

Despite the low initial PPVs, the retrained models in DeepVariant separate errors from real variants with high accuracy in the WGS datasets (PPVs of 99.0% and 97.3% for SOLID and PacBio, respectively), though with a larger loss in sensitivity (candidates 82.5% and final 76.6% for SOLID and 93.4% and 88.5% for PacBio) than other technologies. Additionally, despite the challenges of retraining deep learning models with limited data, the exome datasets also perform strikingly well, with a small reduction in sensitivity (from 91.9% to 89.3% and 94.0% to 92.6% for Ion and TruSeq candidates and final calls) for a substantial boost in PPV (from 8.1% to 99.7% and 65.3% to 99.3% for Ion and TruSeq). The performance of DeepVariant compares favorably to those of callsets submitted to the Genome in a Bottle project site using tools developed specifically for each NGS technology and to callsets produced by the GATK or samtools (Table S7).

The accuracy numbers presented here shouldn’t be viewed as the maximum achievable by either the sequencing technology or DeepVariant. For consistency, we used the same model architecture, image representation, training parameters, and candidate variant criteria for each technology. Because DeepVariant achieves high PPVs for all technologies, the overall accuracy (F1), which is the harmonic mean of sensitivity and PPV, is effectively driven by the sensitivity of the candidate callset. Improvements to the data processing steps before DeepVariant and the algorithm used to identify candidate variants will likely translate into substantial improvements in overall accuracy, particularly for multi-allelic indels. Conversely, despite its effectiveness, representing variant calls as images and applying general image-classification models is certainly suboptimal, as we were unable to effectively encode all of the available information in the reads and reference into the three-channel image.

Taken together, our results demonstrate that the deep learning approach employed by DeepVariant is able to learn a statistical model describing the relationship between the experimentally observed NGS reads and genetic variants in that data for potentially any sequencing technology. Technologies like DeepVariant change the problem of calling variants from a laborious process of expert-driven, technology-specific statistical modeling to a more automated process of optimizing a general model against data. With DeepVariant, creating a NGS caller for a new sequencing technology becomes a simpler matter of developing the appropriate preprocessing steps, training a deep learning model on sequencing data from samples with ground truth data, and applying this model to new, even non-human, samples.

At its core, DeepVariant (1) generates candidate entities with high sensitivity but low specificity, (2) represents the experimental data about each entity in a machine-learning compatible format and then (3) applies deep learning to assign meaningful biological labels to these entities. This general framework for inferring biological entities from raw, errorful, indirect experimental data is likely applicable to other high-throughput instruments.

## Acknowledgements

We would like to thank Justin Zook and his collaborators at NIST for their work developing the Genome in a Bottle resources, the Verily sequencing facility for running the NA12878 replicates, and our colleagues at Verily and Google for their feedback on this manuscript and the project in general.

## Limitations

This manuscript documents the work on DeepVariant leading up to the precisionFDA Truth Challenge in May 2016 in which DeepVariant won the “Highest SNP Performance” award. Since then, at the request of the scientific community, we have entirely rewritten DeepVariant from scratch in order to make it available as open source software. As a result, several improvements to the DeepVariant method aren’t captured in the analyses presented here including switching to TensorFlow^33^ to train the model, using the inception_v3 neural network architecture, and using a multichannel tensor representation for the genomics data instead of an RGB image. The latest results and DeepVariant code are available on GitHub (https://github.com/google/deepvariant/) and additional benchmarking analyses are available from a variety of independent third parties.

## Online methods

### Haplotype-aware realignment of reads

Mapped reads are preprocessed using an error-tolerant, local De-Bruijn-graph-based read assembly procedure which realigns them according to their most likely derived haplotype. Candidate windows across the genome are selected for reassembly by looking for any evidence of possible genetic variation such as mismatching or soft clipped bases. The selection criteria for a candidate window are very permissive so that true variation is unlikely to be missed. All candidate windows across the genome are considered independently. De-Bruijn graphs are constructed using multiple fixed k-mer sizes (from 20 to 75, inclusive, with increments of 5) out of the reference genome bases for the candidate window as well as all overlapping reads. Edges are given a weight determined by how many times they are observed in the reads. We trim any edges with weight less than three, except edges found in the reference are never trimmed. Candidate haplotypes are generated by traversing the assembly graphs and the top two most likely haplotypes are selected which best explain the read evidence. The likelihood function used to score haplotypes is a traditional pair HMM with fixed parameters that do not depend on base quality scores. This likelihood function assumes that each read is independent. Finally, each read is then realigned to its most likely haplotype using a Smith-Waterman-like algorithm with an additional affine gap penalty score for homopolymer indels. This procedure updates both the position and the CIGAR string for each read.

### Finding candidate variants

Candidate variants for evaluation with the deep learning model are identified with the following algorithm. We consider each position in the reference genome independently. For each site in the genome we collect all the reads that overlap that site. The CIGAR string of each read is decoded and the corresponding allele aligned to that site is determined, which are classified into either a reference-matching base, a reference-mismatching base, an insertion with a specific sequence, or a deletion with a specific length. We count the number of occurrences of each distinct allele across all reads. An allele is considered a candidate if it satisfies:

~~~
def is_candidate(counts, allele):
  allele_count = counts[allele]
  total_counts = sum(counts.values())
  return not is_reference_base(allele)
     and allele_count >= min_count
     and allele_count / total_count >= min_fraction
~~~

If any candidates pass our calling thresholds at a site in the genome, we emit a VCF-like record with chromosome, start, reference bases and alternate bases, where reference bases and alternate bases are the VCF-compatible representation of all of the passing alleles.

We filter away any unusable reads (see is_usable_read() below) if it is marked as a duplicate, as failing vendor quality checks, isn’t aligned or if this isn’t the primary alignment, mapping quality is less than 10, or the read is paired and not marked as properly placed. We further only include read bases as potential alleles if all of the bases in the alleles have a base quality >= 10. We only emit variant calls at standard (ACGT) bases in the reference genome. It is possible to force candidate variants to be emitted (randomly with probability of p) at sites with no alternate alleles, which are used homozygous reference training sites. There’s no constraint on the size of indels emitted, so long as the exact position and bases are present in the cigar string and they are consistent across multiple reads.

### Creating images around candidate variants

The second phase of DeepVariant encodes the reference and read support for each candidate variant into an RGB image. The pseudo-code for this component is shown below; it contains all of the key operations to build the image, leaving out for clarity error handling, code to deal with edge cases like when variants occur close to the start or end of the chromosome, and the implementation of non-essential and/or obvious functions.

~~~
WIDTH = 221
HEIGHT = 100;
def create_pileup_images(candidate_variants):
  for candidate in candidate_variants:
   for biallelic_variant in split_into_biallelics(candidate):
     start = biallelic_variant.start - (WIDTH-1) / 2
     end = WIDTH - span_start
     ref_bases = reference.get_bases(start, end)
     image = Image(WIDTH, HEIGHT)
     row_i = fill_reference_pixels(ref, image)
     for read in reads.get_overlapping(start, end):
       if row_i < HEIGHT and is_usable_read(read):
         add_read(image, read, row_i)
         row_i += 1
     yield image

def fill_reference_pixels(ref, image):
   for row in range(5):
    for col in range(WIDTH):
       alpha = 0.4
       ref_base = ref[col]
       red = get_base_color(ref_base)
       green = get_quality_color(60) # The reference is high quality
       blue = get_strand_color(True) # The reference is on the positive strand
       image[row, col] = make_pixel(red, green, blue, alpha)
   return 5

def add_read(image, read, row_i):
# Don’t incorporate reads with a low quality base at the call position. This
# function still returns true because the image isn’t yet full.
# base_quality_at_call_position() returns the quality of the base aligned to
# our call.start, or 255 if no bases are aligned there.
if base_quality_at_call_position(read) < MINIMUM_BASE_QUALITY:
  return

for ref_pos, read_pos, cigar_elt in per_base_alignment(ref, read):
  read_base = None
  if cigar_elt in {‘D’, ‘I’}:
    col = ref_pos - 1
    read_base = INDEL_ANCHORING_BASE
  elif cigar_elt == ‘M’:
    col = ref_pos
    read_base = read.bases[read_pos]
  if read_base:
    quality = min(read.quals[read_pos], read.mapping_quality)
    alpha = get_base_alpha(read_base, ref[col], read, call)
    red = get_base_color(read_base)
    green = get_quality_color(quality)
    blue = get_strand_color(read.is_on_positive_strand)
    image[row_i, col] = make_pixel(red, green, blue, alpha)
def make_pixel(red, green, blue, alpha):
    return RGB(int(alpha * red), int(alpha * green), int(alpha * blue))
def get_base_alpha(read_base, ref_base, read, call):
  # read_supports_alt_allele() returns True if the read supports the alt_allele.
  # This is implemented by associating each alternative allele in our candidate
  # variants with a list of the names of the reads that contained that allele.
  alpha1 = 1.0 if read_supports_alt_allele(read, call.alt_allele) else 0.6
  alpha2 = 0.2 if read_base == ref_base else 1.0
  return alpha1 * alpha2
def get_base_color(base):
  base_to_color = {‘A’: 250, ‘G’: 180, ‘T’: 100, ‘C’: 30}
  return base_to_color.get(base, 0)
def get_quality_color(quality):
  return int(254.0 * (min(40, quality) / 40.0))
def get_strand_color(on_positive_strand):
  return 70 if on_positive_strand else 240
def is_usable_read(read):
  return (read.has_alignment and
          not (read.is_duplicate or read.failed_vendor_quality_checks or
               read.is_secondary or read.is_supplementary) and
          (not read.is_paired or read.is_properly_placed) and
          read.mapping_quality >= 10)
~~~

The actual implementation of this code uses a reservoir sampler to randomly remove reads at locations where there’s excessive coverage. This downsampling occurs conceptually within the reads.get_overlapping() function but occurs in our implementation anywhere where there’s more than 10,000 reads in a tiling of 300 bp intervals on the chromosome.

### Deep learning

DistBelief^34^ was used to represent models, train models on labeled images, export trained models, and evaluate trained models on unlabeled images. We adapted the inception v2 architecture to our input images and our three-state (hom-ref, het, hom-alt) genotype classification problem. Specifically, we created an input image layer that rescales our input images to 299 × 299 pixels without shifting or scaling of our pixel values. This input layer is attached to the ConvNetJuly2015v2^5^ CNN with 9 partitions and weight decay of 0.00004. The final output layer of the CNN is a three-class Softmax layer with fully-connected inputs to the preceding layer initialized with Gaussian random weights and stddev of 0.001 and a weight decay of 0.00004.

The CNN was trained using stochastic gradient descent in batches of 32 images with 8 replicated models and RMS decay of 0.9. For the the Platinum Genomes, precisionFDA, NA12878 replicates, mouse and genome build experiments multiple models were trained (using the product of learning rates of [0.00095, 0.001, 0.0015] and momenta [0.8, 0.85, 0.9]) for 80 hrs or until training accuracy converged, and the model with the highest accuracy on the training set selected as the final model. For the multiple sequencing technologies experiment, a single model was trained with learning rate 0.0015 and momentum 0.8 for 250,000 update steps. In all experiments unless otherwise noted the CNN was initialized with weights from the imagenet model ConvNetJuly2015v2^5^.

### DeepVariant inference client and allele merging

At inference time each biallelic candidate variant site represented as a pileup image is presented as input to the trained CNN. After a forward pass through the network a three-state probability distribution is returned. These probabilities correspond to the biallelic genotype likelihood states of {P(homozygous reference), P(heterozygous), P(homozygous variant)} and are encoded directly in the output VCF record as the phred scaled GL field. Variant calls are emitted for all sites where the most likely genotype is either het or hom-alt with at least a Q4 genotype confidence. Finally all biallelic records at the same starting position are merged into multiallelic records to facilitate comparisons with other datasets.

### Genome in a Bottle human reference datasets

We used version 3.2.1 of the Genome in a Bottle reference data^35^. We downloaded calls in VCF format and confident called intervals in BED format from:

- NA12878: https://ftp-trace.ncbi.nlm.nih.gov/giab/ftp/release/NA12878_HG001/NISTv3.2.1/
- NA24385: https://ftp-trace.ncbi.nlm.nih.gov/giab/ftp/release/AshkenazimTrio/HG002_NA24385_son/NISTv3.2.1/

The VCF files were converted to Global Alliance for Global Health (GA4GH) protocol buffer format but otherwise were used without further modification.

### Evaluating variant calls

Truth variants and confident reference intervals were parsed from the Genome in a Bottle or other ground standard datasets from the VCF and BED files for their respective samples. Truth variants outside the confident intervals were removed. The evaluation variants were loaded and variants marked as filtered or assigned homozygous reference genotypes were removed. Metrics such as the number of SNPs, number of Indels, insertion / deletion ratio, heterozygous / homozygous non-reference ratio, and transition / transversion ratio (Ti/Tv) were calculated from all remaining evaluation variants.

Evaluation variants were matched to truth variants if they start at the same position on the same chromosome. To compute genotype concordance, we added to the list of matched pairs of evaluation / truth variants all of the unmatched evaluation variants that overlap the confidence intervals with a “virtual” homozygous reference genotype sample. The number of matching genotype is defined as the number of pairs where the genotype alleles of the evaluation variant and truth variant are equal, independent of order. From this we compute the genotyping concordance as:

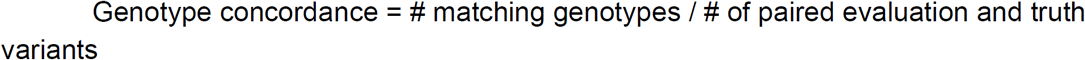

The number of matched pairs is counted as the number of truth positives. Any truth variants without a matched evaluation variant are counted as false negatives. Any unmatched evaluation variants that occur within the confident intervals are counted as false positives. From the number of true positives (TP), false negatives (FN), and false positives (TP) we compute the sensitivity, PPV, and F1 as:

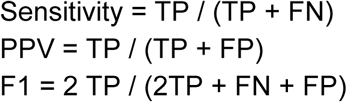

Our evaluation metrics fall between the tolerant hapdip metric^3^ and the strict vcfeval^36^ metrics. In particular, our sensitivity and PPV metrics emphasize discriminating between variant and reference sites, allowing errors in the determination of the exact variant alleles and genotypes. These errors are tallied separately as an allelic error rate and a genotyping error rate. Though we believe this separation is informative and valuable for understanding the types of errors that occur in a variant callset, we appreciate the approaches pursued by other evaluation methods.

### GATK pipeline

For all GATK^6^ analyses (except the Platinum Genomes analysis, see below) we used the Verily production GATK pipeline:

### Versions

~~~
Reference: hg38.genome.fa
dbSNP: v146 on b38 downloaded from NCBI
1000 Genomes Phase 3 callset:
1000G_ALL.wgs.phase3_shapeit2_mvncall_integrated_v5b.20130502.sites.hg38.vcf
downloaded from 1000G FTP
BWA version: 0.7.12
Samtools version: 1.1
Picard version: 2.1.0
GATK version: 3.5
~~~

### BWA

~~~
bwa mem -t 32 fastq1.gz fastq2.gz
    | samtools view -u -
    | samtools sort -@ 12 -O bam -T sorted.bam.sort_tmp -o sorted.bam -
~~~

### Mark Duplicates

~~~
java -Xmx12G -jar picard.jar MarkDuplicates INPUT=sorted.bam
OUTPUT=sorted.deduped.bam ASSUME_SORTED=true CREATE_INDEX=true
MAX_RECORDS_IN_RAM=2000000 METRICS_FILE=MarkDuplicates_metrics.txt
REMOVE_DUPLICATES=false
~~~

After MarkDuplicates, all lanes for the sample are merged into a single BAM file with MergeSamFiles in picard.

### Indel realignment

~~~
java -jar CommandLineGATK_deploy.jar -Xmx4G -R hg38.genome.fa -ip 50 -T
RealignerTargetCreator -I sorted.deduped.merged.bam -known
1000G_ALL.wgs.phase3_shapeit2_mvncall_integrated_v5b.20130502.sites.hg38.vcf -o
realignment_targets.interval_list -nt 8 -mismatch 0.0
java -jar CommandLineGATK_deploy.jar -Xmx4G -R hg38.genome.fa -ip 50 -T
IndelRealigner -I sorted.deduped.merged.bam -targetIntervals
realignment_targets.chr1.interval_list -known
1000G_ALL.wgs.phase3_shapeit2_mvncall_integrated_v5b.20130502.sites.hg38.vcf
--consensusDeterminationModel KNOWNS_ONLY -o sorted.deduped.merged.realigned.bam
~~~

### Base recalibration

~~~
java -jar CommandLineGATK_deploy.jar -Xmx4G -R hg38.genome.fa -T BaseRecalibrator
-I sorted.deduped.merged.realigned.bam -knownSites dbsnp_146.hg38.vcf -o
base_recalibration.table -nct 32 --useOriginalQualities --disable_indel_quals -cov
ReadGroupCovariate -cov QualityScoreCovariate -cov CycleCovariate -cov
ContextCovariate
java -jar CommandLineGATK_deploy.jar -Xmx4G -R hg38.genome.fa -T PrintReads -nct 8
-I sorted.deduped.merged.realigned.bam -BQSR base_recalibration.table
--disable_indel_quals --emit_original_quals -o
sorted.deduped.merged.realigned.recalibrated.bam
~~~

### HaplotypeCaller

~~~
java -jar CommandLineGATK_deploy.jar -Xmx4G -R hg38.genome.fa -ip 50 -T
HaplotypeCaller -I sorted.deduped.merged.realigned.recalibrated.bam -ERC GVCF -o
g.vcf --annotation QualByDepth
java -jar CommandLineGATK_deploy.jar -Xmx4G -R hg38.genome.fa -T GenotypeGVCFs -o
raw_calls.vcf -nt 8 -D dbsnp_146.hg38.vcf --variant g.vcf
~~~

### VQSR

~~~
java -jar CommandLineGATK_deploy.jar -Xmx20G -R hg38.genome.fa -T
VariantRecalibrator --max_attempts 4 -input raw_calls.vcf
-resource:ALL_1000G_phase3,known=false,training=true,truth=true,prior=12.0
1000G_ALL.wgs.phase3_shapeit2_mvncall_integrated_v5b.20130502.sites.hg38.vcf
-resource:dbsnp,known=true,training=false,truth=false,prior=2.0 dbsnp_146.hg38.vcf
-an DP -an QD -an FS -an SOR -an MQ -an MQRankSum -an ReadPosRankSum -mode SNP -nt
4 -tranche 99.5 -recalFile snps.recal -tranchesFile snps.tranches -allPoly
java -jar CommandLineGATK_deploy.jar -Xmx20G -R hg38.genome.fa -T
ApplyRecalibration -input raw_calls.vcf -mode SNP --ts_filter_level 99.5 -recalFile
snps.recal -tranchesFile snps.tranches -o recal.snps.raw.indels.vcf
java -jar CommandLineGATK_deploy.jar -Xmx20G -R hg38.genome.fa -T
VariantRecalibrator --max_attempts 4 -input recal.snps.raw.indels.vcf
-resource:ALL_1000G_phase3,known=false,training=true,truth=true,prior=12.0
1000G_ALL.wgs.phase3_shapeit2_mvncall_integrated_v5b.20130502.sites.hg38.vcf
-resource:dbsnp,known=true,training=false,truth=false,prior=2.0 dbsnp_146.hg38.vcf
-an QD -an DP -an FS -an SOR -an MQRankSum -an ReadPosRankSum -mode INDEL -nt 4
-tranche 99.0 -recalFile indels.recal -tranchesFile indels.tranches -allPoly
java -jar CommandLineGATK_deploy.jar -Xmx20G -R hg38.genome.fa -T
ApplyRecalibration -input recal.snps.raw.indels.vcf -mode INDEL -ts_filter_level
99.0 -recalFile indels.recal -tranchesFile indels.tranches -o final.vcf
~~~

### DeepVariant and GATK on Platinum Genomes NA12878

We trained a deep learning model as described above using only the reads aligned to chromosomes 1 through 18 and evaluated variant calling accuracy on chromosomes 20 to 22 using both our algorithm and the community gold standard GATK best practices pipeline. We reserved chromosome 19 for hyperparameter optimization of the deep learning model. We created a non-overfitted GATK callset in which training does not see the data from chr20-22 by excluding that data during the GATK VQSR step.

For a comparison, we ran GATK v3.3 following Broad best practices as implemented by Google Cloud Genomics + Broad in the alpha version (see https://cloud.google.com/genomics/), run in January 2016 on the NA12878 Platinum Genomes BAM file from https://cloud.google.com/genomics/data/platinum-genomes.

## Supplementary materials

Are available in a separate supplementary materials document.

## Data availability

Most of the data that support the findings of this study are available from Genome in a Bottle or the Mouse Genome project. Some of the data, in particular the 35 NA12878 WGS replicates from the Verily sequencing lab, was licensed from Verily for the current study and so are not publicly available; data are however available from the authors upon reasonable request with permission of Verily.

## Code availability

A production-grade version of DeepVariant was released to GitHub at https://github.com/google/deepvariant. All of the key results and analyses presented here can be reproduced using the open-sourced version of DeepVariant. Custom code was specific to our computing infrastructure and mainly used for simple data analysis tasks.

