## Supplementary Materials for "Creating a universal SNP and small indel variant caller with deep neural networks"

#### Figure 2A details and additional analyses

In both Figure 2A and Figure S1, DeepVariant and GATK calling performance is shown for the Genome in the Bottle benchmark sample NA12878 using 2x101 Illumina HiSeq data from the Platinum Genomes project. The GATK was run in two ways. In the first, GATK best-practices were followed and the variant filtering step (VQSR) was provided data for known variants on both the training and test chromosomes, allowing VQSR to use population variation information to better call variants on the test chromosomes. In the second, we removed all population variation information for the test chromosomes chr20-22, relying on the VQSR model learned only on the training chromosomes, which is more representative of the GATK's calling performance on novel variation. Variants were sorted by QUAL score for DeepVariant and VQSLOD for GATK. Variants that are filtered out in the VCF files are included in the ranking to give a more complete picture of the effectiveness of these ranking methods. This means that the curve includes all candidate variants seen by DeepVariant except those with a homozygous-reference genotype according to the CNN and everything emitted by GATK, including those filtered with LOW_VQSLOD (which, by definition, have a low VQSLOD score).

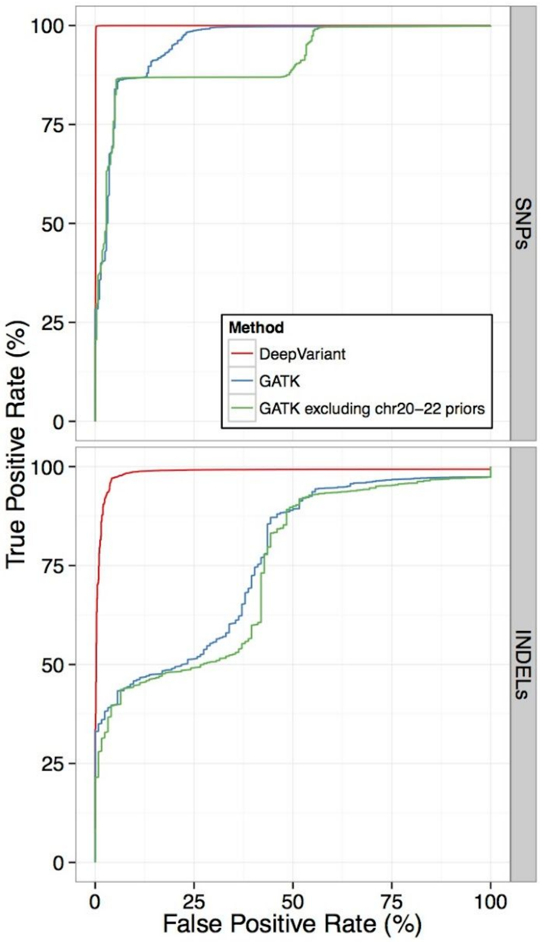


Figure S1: Receiver operating characteristic (ROC) curve for DeepVariant (red) and GATK (green, blue) calls for the Genome in the Bottle benchmark sample NA12878.

Figure 2A and S1 are similar but emphasize different things. The precision-recall plot in Figure 2A gives a better sense of how the end-to-end assay (variant calling) is performing, while the ROC curve in Figure S1 emphasizes the effectiveness of the ranking of true positives relative to false positives, independent of the number of true and false positive variants in each SNP and indel class. In NGS variant calling, a traditional ROC curve can be misleading and is shown here only for completeness. The first of two issues is that the set of false positives is defined as variant calls made into confidently homozygous reference regions by a specific calling method, and so usually differs between calling methods. The second issue is that there is no clear definition of specificity since every allele at every position is a potential true negative. As a consequence, ROC curves across methods are not directly comparable, and so cannot be used to assess the quality of a callset produced by one method relative to another. Precision-recall plot, on the other hand, can be safely compared across methods despite differences in their total number of false positives.

#### DeepVariant vs. GATK on NA12878 replicates

Libraries were prepared from 35 independent replicates of 1ug aliquots of purified genomic DNA isolated from GM12878. During the library preparation process, samples were acoustically sheared to target fragment lengths of 400bp before proceeding through SPRI-based size selection, end repair, a-tailing, adapter ligation, and a final SPRI-based cleanup. The resultant libraries were quantified by Picogreen, Fragment Analyzer, and qPCR. Sequencing was performed using a 2x150 paired-end runs on Illumina HiSeq X sequencers with a targeted sequencing depth of 30x per sample.

Chromosomes 1-18 of the first eight sequenced replicates were used to train a single DeepVariant model by concatenating the labeled pileup images from each replicate into a single training set for DistBelief as previously described. Due to the timing of this experiment, version 2.19 of the Genome in a Bottle reference intervals and variant calls were used to label the genotypes in the training images and to evaluate the quality of the resulting variant calls on held out chromosomes 20-22. The previously described Verily GATK pipeline was used to process each NA12878 sample independently.

### Training and generalization of DeepVariant models across genome builds

DeepVariant was trained on data from human genome builds b37 and applied to b38. 80 hours of training was performed using data from chromosomes chr1-19 of the human NA12878 sample and evaluated on the held out human chromosomes chr20-22 (Table S1). The model trained with read data aligned to b37 of the human reference and applied to b38 data had similar performance (overall F1 = 99.45) to one trained on b38 and then applied to b38 (overall F1 = 99.53) thereby demonstrating the generalizability of the model (Table S1).

Supplementary Table S1: DeepVariant calling across genome builds

| Variants | Training data | Evaluation data | PPV | Sensitivity | F1 |
|---:|---:|---:|:---:|:---:|:---:|
| SNPs + indels | b37 chr1-19 | b38 chr20-22 | 99.93% | 98.98% | 99.45% |
| | b38 chr1-19 | b38 chr20-22 | 99.87% | 99.21% | 99.53% |
| SNPs | b37 chr1-19 | b38 chr20-22 | 99.98% | 99.23% | 99.60% |
| | b38 chr1-19 | b38 chr20-22 | 99.93% | 99.35% | 99.64% |
| Indels | b37 chr1-19 | b38 chr20-22 | 99.60% | 97.35% | 98.46% |
| | b38 chr1-19 | b38 chr20-22 | 99.42% | 98.22% | 98.81% |

### Training and generalization of DeepVariant models across species

In order to evaluate the transferability of a DeepVariant model across species we devised the following experiment. We trained a model using the Platinum Genomes NA12878 read set (aligned to b38) and Genome in a Bottle ground truth labels as described previously. We then applied that model to call variants in the synthetic mouse strain 129S1_SvImJ from the Mouse Genome Project (MGP)[1]. For the sake of comparison we also trained models from the mouse read set using as ground truth the genotypes as provided by MGP.

We downloaded the read files (BAM) and variant calls (VCF) for the synthetic mouse strain 129S1_SvImJ from the MGP website (Table S2).

Supplementary Table S2: Mouse dataset sources

| Sample | Data | Location |
|---|---|---|
| 129S1_SvImJ | BAM | ftp://ftp-mouse.sanger.ac.uk/REL-1502-BAM/129S1_SvImJ.bam |
| | VCF | A combed VCF of ftp://ftp-mouse.sanger.ac.uk/REL-1505-SNPs_Indels/mgp.v5.merged.indels.dbSNP142.normed.vcf.gz and ftp://ftp-mouse.sanger.ac.uk/REL-1505-SNPs_Indels/mgp.v5.merged.snps_all.dbSNP142.vcf.gz |
| | REF | GRCm38 from ftp://ftp-mouse.sanger.ac.uk/ref/GRCm38_68.fa |

The v5 of the mouse callset was created, according to this README, with the following procedure:

> *Reads were aligned to the reference genome (GRCm138) using BWA-MEM v0.7.5-r406 (Li and Durbin, 2009; Li, 2013). Reads were realigned around indels using GATK realignment tool v3.0.0 (McKenna et al., 2010) with default parameters. SNP and indel discovery was performed with the SAMtools v1.1 with parameters:*
>
> *Samtools mpileup -t DP,DV,DP4,SP,DPR,INFO/DPR -E -Q 0 -pm3 -F0.25 –d500*
>
> *and calling was performed with BCFtools call v1.1 with parameters:*
>
> *Bcftools call -mv -f GQ,GP -p 0.99*
>
> *Indels were then left-aligned and normalized using bcftools norm v1.1 with parameters:*
>
> *bcftools norm -D -s -m+indels*
>
> *The vcf-annotate function in the VCFtools package was used to soft-filter the SNP and indel calls. SNP calling was performed for each strain independently. A single list of all polymorphic sites across the genome was then produced from all of the 36 strains' SNP calls. This list was then used to call SNPs again, this time across all 36 strains simultaneously, using the 'samtools mpileup -l' option. The calls from all 36 strains were*

> *then merged into a single VCF file. All strain specific information was retained in the sample columns for each strain. For indels, the same approach was taken with the addition of the indel normalisation step after the initial variant calling. Information regarding the filtering of SNP and indel calls can be found in the VCF file headers in the '##FILTER' and '##source' lines.*

DeepVariant was run using the computational pipeline described above with all default settings. SNP and Indel mutations that were identified in the MGP ground truth set with genotype as 0/0 for this specific mouse were given the hom-ref label and likewise for heterozygous and homozygous variants. No-called sites were ignored during model training.

In order to protect against model overfitting, we divided our human and mouse genomes into a training set of chromosomes and an independent, held-out set of chromosomes (Table S3):

Supplementary Table S3: Training and evaluation chromosomes

| | Training chromosomes | Evaluation chromosomes |
|---|---|---|
| Human NA12878 | chr1-19 | chr20-22 |
| Mouse 129S1_SvImJ | chr1-17 | chr18-19 |

80 hours of training was performed using images prepared from the training chromosomes. After training the model was frozen and applied to call the variants from the read set. The resulting callsets were evaluated on variants on the held out chromosomes only (Table S4). As the Mouse project did not provide confident regions like the Genome in a Bottle project for NA12878, only non-reference variant calls that occur at a site present in the MGP with a genotype of homozygous reference are counted as false positives.

Supplementary Table S4: Calling performance of DeepVariant on human and mouse datasets

| **Variants** | **Training data** | **Evaluation data** | **PPV** | **Sensitivity** | **F1** |
|---|---|---|---|---|---|
| SNPs + indels | Human chr1-19 | Mouse chr18-19 | 99.53% | 97.07% | 98.29% |
| | Mouse chr1-17 | Mouse chr18-19 | 99.90% | 95.85% | 97.84% |
| SNPs | Human chr1-19 | Mouse chr18-19 | 99.98% | 97.86% | 98.91% |
| | Mouse chr1-17 | Mouse chr18-19 | 99.99% | 99.10% | 99.54% |
| Indels | Human chr1-19 | Mouse chr18-19 | 96.41% | 91.75% | 94.02% |
| | Mouse chr1-17 | Mouse chr18-19 | 99.15% | 73.80% | 84.62% |

### DeepVariant training on multiple sequencing technologies

BAM files were downloaded from the Genome in a Bottle project FTP server (Table S5). After downloading the BAM files are fixed up as indicated and converted to GA4GH protocol buffer format for processing with DeepVariant. The conversion preserves all of the essential read information in the BAM.

Supplementary Table S5: Multiple sequence technologies datasets

| Dataset | Sample | BAM FTP location | Notes |
|---|---|---|---|
| TruSeq exome | NA12878 | Nebraska_NA12878_HG001_TruSeq_Exome/NIST-hg001-7001-ready.bam | Exome |
| 10X GemCode 34x WGS | NA12878 | 10XGenomics/NA12878_phased_possorted_bam.bam | Fixed BAM header |
| 10X Chromium 75x WGS | NA12878 | 10Xgenomics_ChromiumGenome/NA12878_GRCh37.bam | Fixed BAM header |
| PacBio raw reads 40x WGS | NA12878 | NA12878_PacBio_MtSinai/sorted_final_merged.bam | Fixed BAM header |
| Ion AmpliSeq exome | NA24385 | ion_exome/HG002_NA24385_SRR1767409_IonXpress_020_rawlib_24038.bam | Exome; Fixed BAM header; Trimmed unneeded BAM tags |
| HiSeq 60x WGS | NA24385 | NIST_HiSeq_HG002_Homogeneity-10953946/NHGRI_Illumina300X_AJtrio_novoalign_bams/HG002.hs37d5.60x.1.bam | |
| HiSeq 31x WGS | NA24385 | NIST_Illumina_2x250bps/novoalign_bams/HG002.hs37d5.2x250.bam | |
| SOLID 85x WGS | NA24385 | NIST_SOLiD5500W/alignment/5500W_HG002_merged.b37.bam | Trimmed unneeded BAM tags |

FTP paths are given relative to:

- For NA12878: ftp://ftp-trace.ncbi.nlm.nih.gov/giab/ftp/data/NA12878/
- For NA24385: ftp://ftp-trace.ncbi.nlm.nih.gov/giab/ftp/data/AshkenazimTrio/HG002_NA24385_son/

Once converted to GA4GH format, candidate variants are identified using the read bases, qualities, QC flags, and mapping information in the original BAM file. The optional local assembly step was skipped for all datasets, as the assembler is tuned for Illumina data. The two exome datasets were trained and evaluated using confident intervals derived from the intersection of the Genome in a Bottle confident intervals and the RefSeq[2] exon intervals.

For training of each dataset, candidate variants were identified using default parameters* as well as emitting reference "variants" at ~0.1% of randomly selected reference bases. Pileup images were created for each candidate variant and labels assigned using Genome in a Bottle truth variants and intervals for the dataset's sample (see methods for details). These labeled images were filtered to only variants occurring on chromosomes 1-19, leaving 20-22 as an independent evaluation set. Training of the deep learning model was carried out for 250,000 steps starting from a model trained against chr1-19 variants from eight NA12878 replicates (see section DeepVariant vs. GATK on NA12878 replicates for details). After training completed the model was frozen and used to evaluate genotype likelihoods as the "technology-trained model".

For evaluation, candidate variants were identified using default parameters* and pileup images were created for each candidate variant on chromosomes 20-22 only. The technology-trained model for the dataset was applied to these images to compute genotype likelihoods and the likelihoods were combined with the candidate variants to create final variant calls (see methods for details). The candidate variants and the final callset were evaluated again using only chr20-22.

*The RAW PacBio read set was called with a slightly different parameter for the minimum fraction required for an alternate indel allele; we require a fraction of 0.18 rather than the default of 0.12 for all other datasets. At 0.12 over ~150M candidates are found, while at 0.18 we only have ~25M variants to consider. Using 0.18 significantly reduces indel sensitivity, from ~60% with 0.12 to around ~40% with 0.18, but is required to make the creation of pileup images tractable in the current implementation. The SNP threshold remains at 0.12 and produces a highly sensitive set at >99%.

#### Comparison of DeepVariant exome calls with technology-specific variant calls submitted to Genome in a Bottle

We sought to compare the quality of DeepVariant calls to baseline callsets for each technology. As we already established the relative performance of DeepVariant and GATK on Illumina WGS data, we focused primarily on non-Illumina WGS and exome datasets. The challenge is that each technology uses a different data processing pipeline needing dataset-specific settings that are often not documented to produce optimal results. Therefore, we first sought SNP and indel variant calls submitted to Genome in a Bottle by the read data depositors as these are likely already optimized for calling performance on that technology. If not available, we applied the Verily GATK pipeline or, when that proved impossible, samtools, as an alternative variant calling option. The dataset and comparison callsets are given in Table S6.

It's important to recognize that these are apples-to-oranges comparisons. There is no way to ensure information on our evaluation chromosomes were not used to tune the submitters calling pipelines. Given that many tools, like the GATK, make direct use of population variation information to aid in filtering variants, we should expect these callsets to be biased towards higher quality calls. Additionally, in some cases the submitters have used more information than DeepVariant to make calls, such as Ion AmpliSeq exome calls which used four lanes of data rather than our single lane. Finally, the callsets can differ in what regions of the genome were called, an acute issue for the exome datasets. To mitigate differences in exome intervals, we further intersected our RefSeq intervals down to those overlapping the calling intervals provides for the two exome datasets. Nevertheless, we feel that these issues are outweighed by the value of natural comparison points to assess the effectiveness of DeepVariant on these technologies.

Supplemental Table S6: Comparison datasets for Genome in a Bottle analysis

| Dataset | Comparator callset | Comparator notes |
|---|---|---|
| TruSeq exome | ftp://ftp-trace.ncbi.nlm.nih.gov/giab/ftp/data/NA12878/Nebraska_NA12878_HG001_TruSeq_Exome/NIST-hg001-7001-ensemble.vcf and GATK | An ensemble callset that includes calls from the GATK HaplotypeCaller, UnifiedGenotyper, and FreeBayes over the TruSeq exome targeted regions (BED). |
| 10X GemCode 34x WGS | None | No callset submitted to Genome in a Bottle. Focusing on Chromium callset from 10x instead. |
| 10X Chromium 75x WGS | ftp://ftp-trace.ncbi.nlm.nih.gov/giab/ftp/data/NA12878/10Xgenomics_ChromiumGenome_LongRanger2.1_09302016/NA12878_hg19/NA12878_hg19_phased_variants.vcf.gz and GATK | |
| PacBio raw reads 40x WGS | Samtools; we could not get GATK to run on this dataset. | Only structural variant calls were submitted for the Pacific BioSciences WGS data. |
| Ion AmpliSeq exome | ftp://ftp-trace.ncbi.nlm.nih.gov/giab/ftp/data/AshkenazimTrio/analysis/IonTorrent_TVC_03162015/AmpliseqExome.20141120.NA24385.vcf and GATK | variant calls from the Torrent Variant Caller (VCF) made on the Ion effective intervals (BED). The TVC caller used all four lanes of Ion torrent exome data, but DeepVariant made its call only one a single lane of data. |
| HiSeq 60x WGS | None | Not analyzed as DeepVariant performance already well-established on Illumina data |
| HiSeq 31x WGS | None | Not analyzed as DeepVariant performance already well-established on Illumina data |
| SOLID 85x WGS | GATK | No calls submitted to Genome in a Bottle for NA24385. There appear to be no maintained variant callers for SOLID data. |

Samtools calling on PacBio raw reads 40x WGS

```
#!/bin/bash

# Only calling on chromosomes 20, 21, and 22.
CHROMS=('20:1-20,000,000' '20:20,000,000-40,000,000' '20:40,000,000-63025520'
'21:1-20,000,000' '21:20,000,000-40,000,000' '21:40,000,000-48129895'
'22:1-20,000,000' '22:20,000,000-40,000,000' '22:40,000,000-51304566')
```

```
# Run calling on each interval separately.
parallel -j ${#CHROMS[@]} "samtools view -u NA12878_PacBio-RAW.bam {} \
  | samtools mpileup -ugf GRCh37.genome.fa - \
  | bcftools call -vmO z -o NA12878_PacBio-RAW.samtools.calls.{}.vcf.gz" :::
${CHROMS[*]}

# Conconcate parallel calling VCFs.
bcftools concat -a -O z --rm-dups all \
 NA12878_PacBio-RAW.samtools.calls.??\:*.vcf.gz \
 -oNA12878_PacBio-RAW.samtools.calls.vcf.gz
tabix NA12878_PacBio-RAW.samtools.calls.vcf.gz

# Filter recommendations taken from bcftools website with depth of 40x.
bcftools filter -O z -o NA12878_PacBio-RAW.samtools.calls.filtered.vcf.gz \
  -s FAIL -i'DP < 67 && QUAL > 10 & DP >= 3' --SnpGap 3 \
  NA12878_PacBio-RAW.samtools.calls.vcf.gz
tabix NA12878_PacBio-RAW.samtools.calls.filtered.vcf.gz
```

Table S7 shows the PPV, sensitivity, and F1 metric of the DeepVariant and comparator callsets on the previously-indicated regions on the held-out chromosomes 20-22. For exomes the evaluation interval is the intersection of the targeted regions with the RefSeq intervals on chromosomes 20-22.

Supplementary Table S7: Comparison of technology specific callsets and DeepVariant for SNPs + indels combined

| Data | Caller | Sensitivity | PPV | F1 |
|---:|---:|---|---|---|
| Ion AmpliSeq exome | DeepVariant | 94.12% | 99.79% | 96.87% |
| | TVC | 96.47% | 98.11% | 97.28% |
| | GATK | 93.24% | 19.15% | 31.78% |
| Illumina TruSeq exome | DeepVariant | 93.01% | 99.39% | 96.09% |
| | Ensemble | 92.92% | 98.08% | 95.43% |
| | GATK | 91.02% | 99.30% | 94.98% |
| 10X Chromium 75x WGS | DeepVariant | 98.73% | 99.91% | 99.32% |
| | Long-ranger | 98.13% | 98.26% | 98.19% |
| | GATK | 99.08% | 94.62% | 96.80% |
| PacBio raw reads 40x WGS | DeepVariant | 88.51% | 97.25% | 92.67% |
| | samtools | 89.34% | 40.89% | 56.10% |
| SOLID 85x | DeepVariant | 76.62% | 99.01% | 86.39% |

| | | | | |
|---|---|---|---|---|
| | GATK | 73.91% | 84.26% | 78.75% |

As noted in the "evaluation of variants" section, the difference between evaluation methods may be exaggerated in this multiple sequencing technologies experiment. We intentionally chose to use the already aligned BAM files as provided by Genome in a Bottle in order to highlight the robustness of DeepVariant to variation in input alignments without applying local assembly, which may perform better on Illumina read than other NGS read types. One consequence of this choice, though, is that DeepVariant will only call alleles present in the CIGAR elements of the BAMs, which vary in their accuracy depending on the sophistication of the aligner and post-alignment cleanup steps performed during processing by each technology's data depositor. The DeepVariant CNN is sufficiently robust to train accurate genotyping models even with errorful allele determination, as evident by the high PPV values, but inevitably produces variant calls with incorrect alleles at any site where the reads have been aligned with an incorrect allele in their CIGAR elements. As noted in the main text, better pre-processing via tools like the GATK's IndelRealigner[3] or technology-agnostic local assembly will improve the alleles emitted by DeepVariant. Additionally, it is possible that our comparator callsets may use variant representations that are differentially penalized by our evaluation tool. Because of these concerns, we ran both our internal evaluation tool and vcfeval (version 3.6.2) and note that the results are quite concordant between both methods. The full output is available as a supplementary datafile.

#### General guidance on selection of training data for DeepVariant

For most of the experiments presented here DeepVariant was trained on 8 whole genome replicates of NA12878 sequenced under a variety of conditions related to library preparation. These conditions include loading concentration, library size selection, and laboratory technician. We believe this diversity in training data is one reason that DeepVariant is able to generalize to a variety of new datasets. In general in order to be robust to a particular class of errors training data sequenced from that class should be included in the set of training data. We also believe that by training with even more sequencing data will improve the performance of DeepVariant even further but there is a tradeoff with the number of training iterations required as the number of training examples increases.
